# Comparative effects of inactivation and dopamine receptor agents in the dorsolateral and dorsomedial striatum on performance of action sequences in rats

**DOI:** 10.64898/2026.01.22.701165

**Authors:** Karly M. Turner, Anna Svegborn, Trevor W. Robbins

## Abstract

**Rationale:** Recent research on habits and skills has produced a wave of new theories regarding the shift in control from medial to lateral regions of the dorsal striatum, and how these regions are implicated in the selected and executed of action sequences.

**Objectives:** To examine the comparative effects of muscimol/baclofen inactivation and dopamine D1 and D2 receptor agents in the dorsomedial (DMS) and dorsolateral (DLS) striatum on the performance of skilled action sequences.

**Methods:** Infusions were made in well-trained rats using the five-step nose poke task to isolate the effects on initiation, execution and termination components of skilled action sequences.

**Results:** DLS inactivation produced sequencing deficits like those observed with pre-training lesions, indicating that the DLS is critical for both the acquisition and performance of sequences. Behaviour was unchanged following DMS inactivation, consistent with models of DMS disengagement following training. Infusions of D1 and D2 antagonists did not alter behaviour, however the D2 receptor agonist quinpirole increased sequence errors at a low dose and reduced sequences at the high dose in the DLS. DLS manipulations impaired sequence initiation and termination as well as reward transitions, while the ‘chunking’ ballistic response pattern was largely unaltered, indicating that between-but not within-sequence actions rely on the DLS.

**Conclusions:** Skilled action sequencing, including ‘chunk’ transitions was dependent on DLS and its modulation by D2 receptors, but not on DMS function. Using a novel sequencing task, these results support the dissociable and dopamine-dependent role of the dorsal striatum subregions in performing skilled motor actions.

## Introduction

The ability to efficiently perform routine action sequences plays a vital function in our daily lives. Actions must be learned and then executed in the appropriate order to achieve the desired outcome. With practice, sequences are performed fluidly and with minimal effort. This is thought to occur through the binding of individual actions into ‘chunks’ to allow rapid and refined, albeit rigid, motor patterns to be executed with minimal cognitive effort (Graybiel 1998; Ostlund et al. 2009). This trajectory towards automaticity is common to skills, habits and chunked action sequences. Distinguishing each of these terms and how they interact has been reviewed extensively and additional criteria are required to isolate clear definitions (Dezfouli et al. 2014; Du et al. 2022; Garr and Delamater 2019; Graybiel and Grafton 2015; Robbins and Costa 2017; Song and Lee 2024). For example, habits are identified as insensitivity to outcome devaluation and contingency degradation (Garr and Delamater 2019; Watson et al. 2022). While skills are less clearly defined, they typically feature the optimisation of motor action selection and execution leading to greater speed and accuracy with little variability (Yadav and Duque 2023). Impairments in the development of skills and/or dysfunctional habit formation is apparent in a range of neuropsychiatric disorders, including Obsessive-Compulsive Disorder (OCD) (Gillan et al. 2014; Graybiel and Rauch 2000), addiction (Belin et al. 2009; Everitt and Robbins 2016), Parkinson’s Disease (Cohen and Pourcher 2007; Ruitenberg et al. 2015) and Huntington’s Disease (Heindel et al. 1988; Heindel et al. 1989). Moreover, many of these conditions are associated with altered striatal functionality. Therefore, investigating adaptations within striatal circuits as new skills are mastered will improve our understanding of healthy and dysfunctional brain processes.

Within the dorsal striatum, the segregation of medial and lateral subregions provides further functional precision. The dorsomedial striatum (DMS) is central to learning and goal-directed behaviours, whereas the dorsolateral striatum (DLS) is critical for the development of habits and skills (Balleine and O’Doherty 2010; Graybiel 2008; Yin and Knowlton 2006). This has been demonstrated using various behavioural paradigms across species, suggesting functional homology of these regions is reasonably well conserved (Balleine and O’Doherty 2010). Yet, how the balance of control shifts as an action becomes more habitual or skilful has remained elusive. Various studies have demonstrated that both regions are engaged in task-related activity to varying degrees across learning and performance (Bassett et al. 2015; Graybiel 2008; Thorn et al. 2010). Therefore, both regions appear to be dynamically adapting across stages of habit or skill formation, just as the balance of psychological processes controlling action gradually shifts (Dolan and Dayan 2013).

Sub-regions of the dorsal striatum retain the capacity to act independently as demonstrated by ablating or silencing the alternative region. Disrupting DMS function induces habitual behaviour even very early in learning, and DLS disruption leads to goal-directed behaviour in extensively trained animals that would otherwise show habits (Killcross and Coutureau 2003; Yin et al. 2004). Given the DMS and DLS are independently capable of controlling behaviour, yet appear to be simultaneously active, it has been suggested that the shift from goal-directed to habitual behaviours, and separately the shift towards skilled execution of actions, is co-ordinated by parallel or competitive processing between these striatal sub-regions (Balleine et al. 2009; Belin et al. 2009; Kupferschmidt et al. 2017; Thorn et al. 2010; Turner et al. 2022).

Similar models of competitive and parallel processing exist for the two major outputs of the basal ganglia, the direct and indirect pathways (Cui et al. 2013). These pathways coincide with the expression of dopamine D1 receptors on the direct pathway neurons (dSPNs) and D2 receptors on the indirect pathway neuron (iSPNs), with only ∼5% of neurons co-expressing both receptor types (Bertran-Gonzalez et al. 2008). It was originally proposed that these pathways operate as a ‘Go’ and ‘Nogo’ pathway respectively, consistent with reports that D1-R activation increases locomotion and D2-R activation reduces movement (Kravitz et al. 2010). However, more recent hypotheses propose D1 dSPNs activity is important for triggering specific actions with D2 iSPNs inhibiting competing actions such that only the selected action is executed (Bariselli et al. 2019; Cui et al. 2013; Fang and Creed 2024; Kravitz et al. 2010). This hypothesis has been refined further, such that D2 iSPNs provide highly selective inhibition, rather than blanket inhibition of all other actions (Klaus et al. 2017). It is important to note that D2 receptors have 10- to 100-fold higher affinity for dopamine than D1 receptors, so under conditions of low concentrations of dopamine (i.e., tonic signalling) there will be greater activation of the D2 indirect pathway, however high concentrations (i.e., phasic release) will activate the D1 direct pathway (Gerfen 2022; Klein et al. 2019; Missale et al. 1998; Tritsch and Sabatini 2012). Aside from the different downstream pathways activated following D1-R and D2-R activation, there are also local collaterals between SPNs within the striatum, allowing D2-expressing iSPNs to inhibit activity of D1-expressing dSPNs (Matamales et al. 2020; Taverna et al. 2008). Therefore, phasic dopamine will not just activate D1 dSPNs, but with relatively less D2 activation reduced collateral inhibition of D1 dSPNs will occur, further increasing likelihood of behaviour initiation. Whereas lower levels of dopamine will lead to relatively greater D2-R activation and ultimately action termination (Cui et al. 2013; Geddes et al. 2018). Therefore, coordination of these systems is critical for the selection and execution of specific action sequences.

In this study we examined the role of the DMS and DLS in the performance of a heterogenous action sequence before isolating the role of D1 and D2 receptors within each subregion. We first examined how action sequencing was altered by local DMS and DLS inactivation using standard methods based on microinfusion of a mixture of the GABA-A agonist muscimol and the GABA-B agonist baclofen, before testing effects of intra-striatal dopamine receptor agents within each subregion (the D2-R antagonist raclopride, D2-R agonist quinpirole and the D1-R antagonist SCH23390). After extensive training, it was predicted that the DLS would be required for optimal task performance, however the DMS would no longer play a critical role. Consistent with the selective ‘go’ and ‘nogo’ framework, it was predicted that D2 manipulations would impact action selection, with more errors and slower initiation and termination elements, whereas D1 manipulations were expected to impact the speed of action execution.

## Methods & Materials

### Animals

Adult male Lister-hooded rats (280-300g; Charles River, UK) were housed within a temperature (21°C) and humidity-controlled environment in open top cages with aspen bedding and a housing tube on reversed 12-h light cycle (lights off 07:00) in groups of four. Rats were food-restricted (>90% of free-feeding weight) with free access to water and given exposure to reward pellets. All procedures were conducted in accordance with the United Kingdom Animal (Scientific Procedures) Act of 1986 and were approved by local ethical review at the University of Cambridge.

### Apparatus

Testing was conducted in operant chambers (Campden Instruments, UK) with five nose poke receptacles arranged in a horizontal array and a reward receptacle connected to a pellet dispenser on the opposing wall (Robbins 2002). A house light was mounted on the ceiling, and the chamber was contained within a sound attenuated box. Cameras (SpyCameraCCTV, UK) were mounted above the chambers to record behaviour remotely. A combination of custom and WhiskerServer software was used to operate the chambers and record responses (Cardinal and Aitken 2010).

### Sequential Nose poke Task

Naïve rats were trained to perform the 5-step sequential nose poke task (SNT) as described in (Turner et al. 2022). Briefly, rats were habituated to the chambers then trained to make nose poke responses. To avoid biasing responding to any hole, a unique training protocol was used where the five nose poke holes would illuminate one at a time from left to right before the reward receptacle was illuminated and a sucrose pellet was delivered (AIN76A, 45mg; TestDiet, UK). However, the time taken to move from one hole to the next incremented with each trial, delaying reward delivery. If a nose poke was detected while the hole was illuminated, it would immediately progress to the next hole, thereby increasing the rate of reinforcement. Rats quickly learned to nose poke the illuminated hole and developed the left to right sequence. Once they had performed at least 15 whole sequences (1-2-3-4-5) within a 30 min session, the illuminated hole would only progress after a nose poke was detected. Once they performed 50 sequences at this stage, the holes were no longer illuminated, and the rats needed to produce the 5-step sequence without cues. At this point, if an error was made, the program waited for the correct response and then progressed (e.g., 1-2-**4**-3-4-5-reward). Upon reaching the criteria of >50 sequences within a session, they moved to the final stage where incorrect nose pokes were punished by a 5s time out and the rat needed to start back at hole 1 on the next sequence. Training on this stage was conducted for a minimum of 15 sessions.

### Surgery

After extensive SNT training, rats underwent surgery to implant cannula into the DMS and DLS in performance matched groups. Stereotaxic surgery was conducted under 2-3% isoflurane anaesthesia with a local injection of bupivacaine (2mg/kg s.c. at 0.8ml/kg; Sigma) and systemic antibiotic (10mg/kg s.c. at 1ml/kg; Baytril). Cannula (22G; PlasticsOne) were implanted bilaterally in the DMS (AP -0.4, ML ±2.2, DV -4.0) or DLS (AP +0.7, ML ±3.6, DV -4.5) and secured with four anchoring screws and dental cement. Rats were treated with Metacam (1mg/kg; Boehringer Ingelheim) pre- and three days post-operatively and rehoused in groups of four. After at least seven days recovery, rats were food restricted and returned to daily SNT sessions until returning to baseline performance levels.

### Infusions

A 10ml Hamilton syringe and pump were used to administer 0.5µl per hemisphere of each drug at 0.3ml/min flow rate (801N, Hamilton, US) via injectors with +0.5mm projection from the guide cannula (28G, PlasticsOne). The awake rat was gently restrained, dummy cannula removed, and injectors inserted into the guide cannula. Injectors were left in place for 1 min then the infusion was started. The injectors were left in place for an additional minute after the infusion to allow diffusion of the fluid before removal and the reinsertion of the dummy cannula. Rats were returned to their home cage to rest before being transported to the operant chambers to begin testing 10min after the completion of the infusion. To habituate rats to the infusion procedure, injectors were first inserted but with no infusion and repeated the next day with a single saline infusion. Rats were retrained in between each infusion session to restore baseline performance levels.

### Drugs

Infusions were made using muscimol hydrobromide (Sigma, USA) and baclofen hydrochloride (Sigma, USA) (1:1 of 1.0mM baclofen and 0.1mM muscimol; MB), SCH23390 (Sigma, USA; 0.5mg/hemisphere, 1.0mg/hemisphere), raclopride (Tocris Bioscience, UK; 0.05mg/hemisphere, 0.5mg/hemisphere) and quinpirole (Sigma, USA; 0.1mg/hemisphere, 1.0mg/hemisphere). All drugs were dissolved in saline and stored at -20°C until the day of use.

### Histology

Rats were perfused (0.01M PBS with 12.5g sodium nitrite for every 2.5L PBS, then 4% PFA in PBS) and brain removed for storage in 4% paraformaldehyde at room temperature overnight on a shaker. They were then rinsed in phosphate buffered saline and transferred to 30% sucrose solution and kept at 4°C until they sank. The brains were then rapidly frozen and sectioned on a microtome (60mm; Leica). Sections were then stained with cresyl violet for cannula placement. Cresyl violet working solution contained 0.2M anhydrous sodium acetate, 5M formic acid, 0.5% cresyl fast violet solution (5g cresyl violet in 1L ddH2O and boiled) in ddH20. Sections were mounted on gelatin-coasted slides and dried. Slides were washed through 100% xylene, 50/50% xylene/ethanol, 100% ethanol, water, cresyl working solution, 100% ethanol, differentiated in 100% ethanol with 0.1% acetic acid, then back through 100% ethanol, 50/50% xylene/ethanol and 100% xylene. There were then cover slipped with depex.

### Statistical Analysis

Key measures included sequences initiated, correct and incorrect sequences, sequence duration, reward magazine duration and nose poke duration, as described previously (Turner et al. 2022). Data were analysed using repeated measures or univariate analysis of variance (ANOVA) to compare performance between treatments with post-hoc comparisons or t-tests used to directly compare effects when appropriate (SPSS, IBM). Greenhouse-Geisser corrections were used where the sphericity assumption was violated.

## Results

### Histology

Cannulation placement was determined from Cresyl staining with infusion locations indicated in Figure 1A-C. Rats were excluded if histology revealed misplaced cannula.

**Fig. 1.**
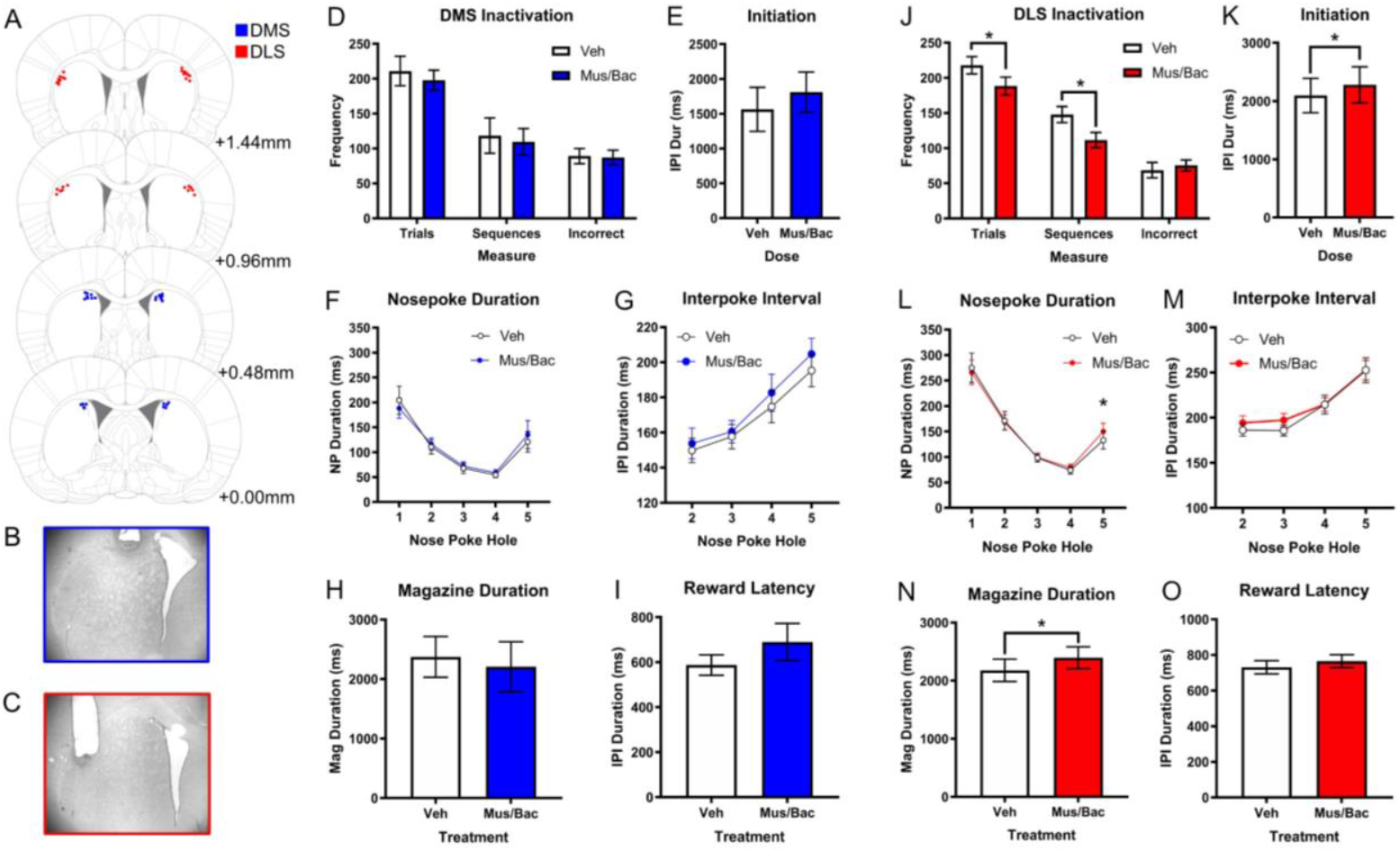
Inactivation of the DMS and DLS during action sequencing. A) Dorsal striatum cannulation placement in the DLS (n=15; red dots) and DMS (n=13; blue dots). B) Histological example of cannula placement in the DMS and C) DLS. D) Infusions of muscimol and baclofen (Mus/Bac) into the DMS did not significantly influence the number of sequences sequencing behaviour, (E) latency to initiate sequences or (F) duration spent in each of the nose poke holes. However, a ballistic response pattern was observed overall. (G) There were no significant effects of inactivation on the interval between nose pokes, (H) time spent in the magazine and (I) a non-significant trend towards longer latency to collect rewards (p=0.07). J). MB Infusions into the DLS significantly reduced the number of trials and correct sequences produced and K) increased the latency to initiate sequences. L) The duration spent in each of the nose poke holes showed a ballistic response pattern with significantly more time spent on the final hole following DLS inactivation. M) There was no significant effect of inactivation on the interval between nose pokes. N) Rats spent significantly longer in the magazine following inactivation but O) latency to collect rewards was not altered. *\*p<0*.*05*

### Inactivation of DMS with muscimol/baclofen mixture

Paired t-tests found no significant effects of DMS inactivation on number of trials, number of correct sequences or incorrect sequences compared to vehicle treatment (Figure 1D). There was no significant effect of inactivation on initiation latency (p=0.35; Figure 1E) or total sequence duration (t_(5)_=-1.67, p=0.16). When comparing the duration of nose pokes on each step of the sequence (NP1-5), there was a significant effect of hole (F_(4,20)_=13.42, p<0.001) but no main effect of infusion (p=0.72) or interaction between hole and infusion (p=0.29; Figure 1F). When comparing the time between each nose poke in the sequence (NP2-5) there was a main effect of hole (F_(3,15)_=17.69, p<0.001) and non-significant trend towards slower intervals between holes following inactivation (p=0.08; Figure 1G) with no interaction (p=0.61). The rewarded magazine duration was also not significantly different after Mus/Bac infusion (p=0.61; Figure 1H). There was a non-significant trend towards a longer reward collection latency (p=0.07; Figure 1I).

### Inactivation of DLS by muscimol/baclofen mixture

Following DLS inactivation, a paired t-test revealed a significant reduction in the number of trials completed (t_(11)_=4.20, p=0.001) and number of sequences generated (t_(11)_=4.16, p=0.002), but not the number of incorrect sequences (t_(11)_=-0.87, p=0.20) compared to vehicle treatment (Figure 1J). There was no significant effect of inactivation on total sequence duration (t_(11)_=-1.65, p=0.13), however the time to initiate trials was significantly longer after MB infusion (t_(11)_=-2.52, p=0.028; Figure 1K). When comparing the duration of nose pokes on each step of the sequence (NP1-5), but there was a significant effect of hole (F_(2,44)_=30.60, p<0.001) and an interaction between hole and infusion (F_(4,44)_=3.11, p=0.024; Figure 1L). Paired t-tests revealed this was due to a longer nose poke duration on hole 5 after MB infusion into the DLS. Although there was a significant effect of hole on the time interval between nose pokes in the sequences (F_(3,33)_=37.09, p<0.001), this measure was not significantly altered by inactivation nor was there an interaction (p’s>0.4; Figure 1M).

A paired t-test found that rats spent significantly longer in the magazine following reward delivery after MB infusion into the DLS (t_(11)_=-2.47, p=0.031; Figure 1N), but reward latency was not prolonged (p=0.11; Figure 1O).

### Effects of Raclopride – D2-R Antagonism in DMS

A repeated measures ANOVA found there was no significant effect of DMS raclopride infusions on the number of trials, correct or incorrect sequences (p’s> 0.2; Figure 2A). There was no significant effect of raclopride on total sequence duration (F_(2,12)_=1.02, p=0.39). When comparing the duration of nose pokes on each step of the sequence (NP1-5), there was a significant effect of hole (F_(1,7)_=11.91, p=0.009) but no main effect of infusion or interaction between hole and infusion (p’s>0.6; Figure 2B). There was a main effect of location on the latency between nose pokes in the sequence (F_(1,6)_=27.67, p=0.001) but no significant effect of dose or interaction (p’s>0.1; Fig 2C). The rewarded magazine nose poke duration and reward latency (p>0.6) was not significantly different after raclopride infusion (Figure 2D-E).

**Fig. 2.**
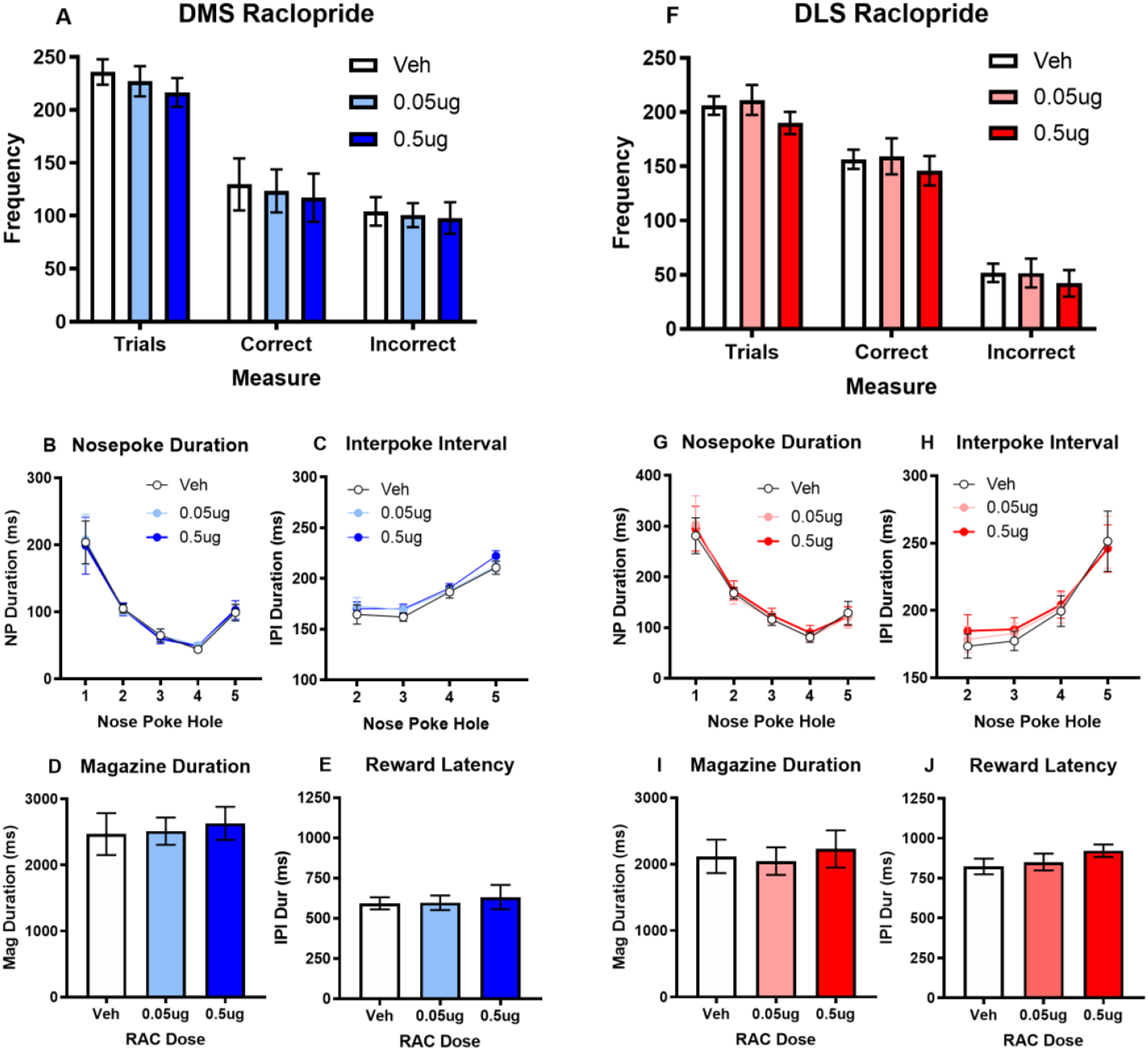
Raclopride infusions in the DMS and DLS. (A, F) Compared to vehicle, infusions of raclopride (RAC) in the DMS and DLS had no effect on any measure of sequencing behaviour, (B, G) nose poke duration, (C, H) interpoke interval or (D, I) magazine duration or (E, J) reward latency. *\*p<0*.*05*

### Effects of Raclopride – D2-R Antagonism in DLS

A repeated measures ANOVA found a significant effect of raclopride infusion into the DLS on the number of trials (F_(2,16)_=3.65, p=0.049; Figure 2F) however the greatest difference was between the 0.05mg and 0.5mg infusions. There were no other significant effects of DLS raclopride infusions on the number of correct sequences or incorrect sequences (p’s>0.5). There was no significant effect of raclopride on total sequence duration (p=0.68). When comparing the duration of nose pokes on each step of the sequence (NP1-5), there was a significant effect of hole (F_(2,12)_=13.31, p=0.001) but no main effect of infusion or interaction between hole and infusion (p’s>0.5; Figure 2G). There was a main effect of location on the latency between nose pokes in the sequence (F_(2,9)_=29.73, p<0.001) but no significant effect of dose or interaction (p’s>0.4; Figure 2H). The rewarded magazine nose poke duration and reward latency (p’s>0.1) was not significantly different after raclopride infusion (Figure 2I-J).

### Effects of Quinpirole – D2-R Agonism in the DMS

A repeated measures ANOVA found quinpirole in the DMS had no significant effect on the number of trials, correct or incorrect sequences (p’s>0.4). There was a non-significant trend towards increased total sequence duration after (low dose) quinpirole (F_(2,14)_=3.56, p=0.056; Figure 3A). When comparing the duration of nose pokes on each step of the sequence (NP1-5), there was a significant effect of hole (F_(2,28)_=17.93, p<0.001) but no main effect of infusion or interaction between hole and infusion (p’s>0.7; Figure 3B). However, the time taken to move between holes varied by hole (F_(2,13)_=11.50, p=0.002; Figure 3C) and was significantly longer after the low dose of quinpirole (F_(2,14)_=5.62, p=0.016; veh vs low dose p=0.029, veh vs high dose p=0.56) without a dose by hole interaction (p=0.22). The rewarded magazine nose poke duration was significantly increased after 0.1mg quinpirole infusion (F_(2,14)_=4.26, p=0.036; Figure 3D) and reward latency (F_(2,14)_=5.56, p=0.017; Figure 3E) was also increased. Therefore, the trend towards longer sequence duration after low dose quinpirole was likely due to a combination of delayed inter-poke interval, reward latency and reward consumption periods.

**Fig. 3.**
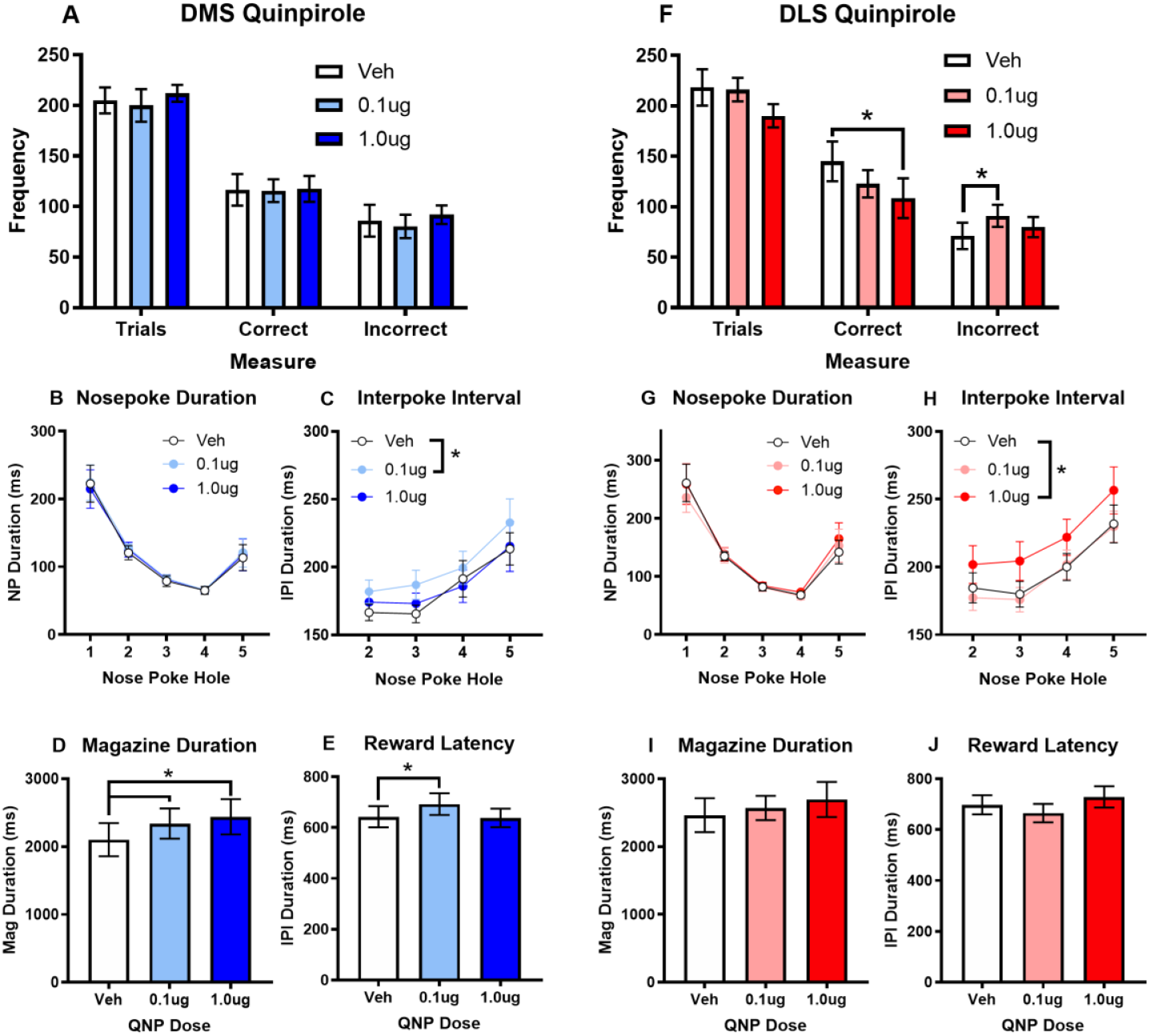
Quinpirole infusions in the DMS and DLS. (A) Quinpirole (QNP) in the DMS had no effect on the number of trials, correct or incorrect sequences, however (F) in the DLS, QNP at the high dose reduced the number of correct sequences and the low dose increased the number of incorrect sequences. (B, G) There was also no change in the time spent in each nose poke hole. However, rats took longer to move between nose poke holes and to the magazine after (D) the low dose in the DMS and (H) high dose in the DLS. (D) DMS time spent in the magazine following a reward was longer after both QNP doses compared to vehicle and (E) reward latency was longer after low dose in the DMS. (I, J) There was no change in the reward duration or latency in the DLS. *\*p<0*.*05*.

### Effects of Quinpirole – D2-R Agonism in the DLS

A repeated measures ANOVA found quinpirole in the DLS significantly reduced the number of correct sequences at the high dose (F_(2,14)_=5.32, p=0.019; veh vs. high dose t_(7)_=2.78, p=0.027) with a non-significant trend towards reduced trials (F_(2,14)_=3.57, p=0.056). The number of incorrect sequences (F_(2,14)_=5.32, p=0.019) was increased, which was driven by an increase in errors after the lower dose of QNP (t_(7)_=-3.64, p=0.008, Figure 3F). There was a non-significant trend towards increased total sequence duration after (high dose) quinpirole (F_(2,14)_=3.39, p=0.063). When comparing the duration of nose pokes on each step of the sequence (NP1-5), there was a significant effect of hole (F_(2,14)_=16.69, p<0.001) but no main effect of infusion (p=0.31) or interaction between hole and infusion (p=0.08, Figure 3G). However, the time taken to move between holes varied by hole (F_(2,11)_=41.53, p<0.001) and was significantly longer after the high dose of quinpirole (F_(2,14)_=13.61, p=0.001; veh vs high dose p=0.006, veh vs low dose p=0.39) without a dose by hole interaction (p=0.68, Figure 3H). The rewarded magazine nose poke duration was not altered (p=0.30; Figure 3I) and the median reward collection latency was significantly longer between doses (F_(2,14)_=3.94, p=0.044, Figure 3J), with neither differing from vehicle treatment. Therefore, sequence duration was extended by the longer inter-poke interval at the higher dose.

### Effects of SCH23390 – D1-R Antagonism in DMS

A repeated measures ANOVA found there was no significant effect of DMS SCH23390 infusions on the number of trials, correct or incorrect sequences (p’s>0.2; Figure 4A). There was also no significant effect on total sequence duration (F_(2,12)_=2.29, p=0.14). When comparing the duration of nose pokes on each step of the sequence (NP1-5), there was a significant effect of hole (F_(1,24)_=12.43, p=0.004) but no main effect of infusion or interaction between hole and infusion (p>0.4; Figure 4B). The latency between nose pokes was significantly different across locations (F_(3,15)_=27.01, p<0.001) and was significantly reduced by low dose of SCH23390 (F_(2,10)_=6.11, p=0.019; veh vs. 1µg p=0.011; Figure 4C) compared to vehicle but without a dose by location interaction (p=0.27). The rewarded magazine nose poke duration and reward latency (p’s>0.1; Figure 4D-E) were not significantly different after SCH23390 infusion.

**Fig. 4.**
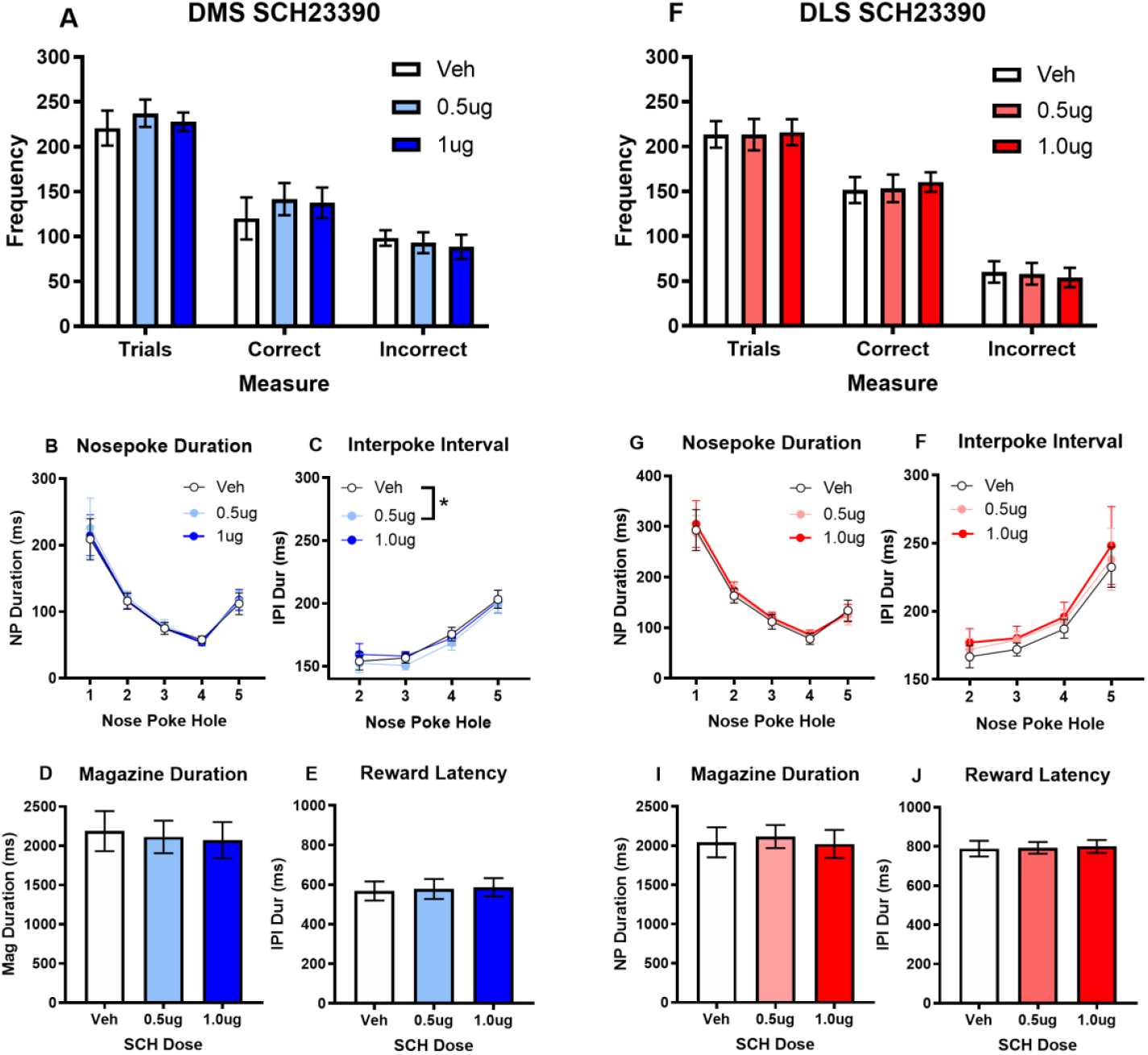
SCH23390 infusions in the DMS and DLS. (A, F) Infusions of SCH23390 (SCH) in the DMS and DLS had no effect on trials, number of sequences or accuracy. (B, G) Although a ballistic response pattern was detected, the time duration spent in each of the nose poke holes was not significantly altered by SCH. (C, F) The interpoke interval (IPI) between nose pokes was significantly longer after low dose SCH in the DMS, but not DLS. (D, I) There was also no significant effect of SCH infusion on time spent in the magazine or (E, J) reward latency. *\*p<0*.*05*

### Effects of SCH23390 – D1-R Antagonism in DLS

A repeated measures ANOVA found no significant effect of DLS SCH23390 infusions on the number of trials, correct or incorrect sequences (p’s = 0.5-0.9; Figure 4F). There was no significant effect of SCH23390 on total sequence duration (F_(1,8)_=0.30, p=0.75). When comparing the duration of nose pokes on each step of the sequence (NP1-5), there was a significant effect of hole (F_(2,32)_=15.55, p=0.001) but no main effect of infusion or interaction between hole and infusion (p’s>0.4; Figure 4G). There was a main effect of location on the latency between nose pokes in the sequence (F_(1,9)_=10.11, p=0.009; Figure 4F) but no significant effect of dose or interaction (p’s>0.1). The rewarded magazine nose poke duration and reward latency was not significantly different after SCH23390 infusion (p’s>0.2; Figure 4I-J).

## Discussion

Acute inactivation of the dorsolateral but not dorsomedial striatum using an intra-striatal muscimol/baclofen mixture produced significant deficits in several parameters of performance in a skilled action sequencing task in rats. By comparison, intra-striatal D1-R or D2-R antagonism had little effect on performance, although intra-striatal infusion of the D2-R agonist quinpirole had significant disruptive effects within both the DMS and DLS, slowing responding and increasing errors (Table I).

**Table I.**
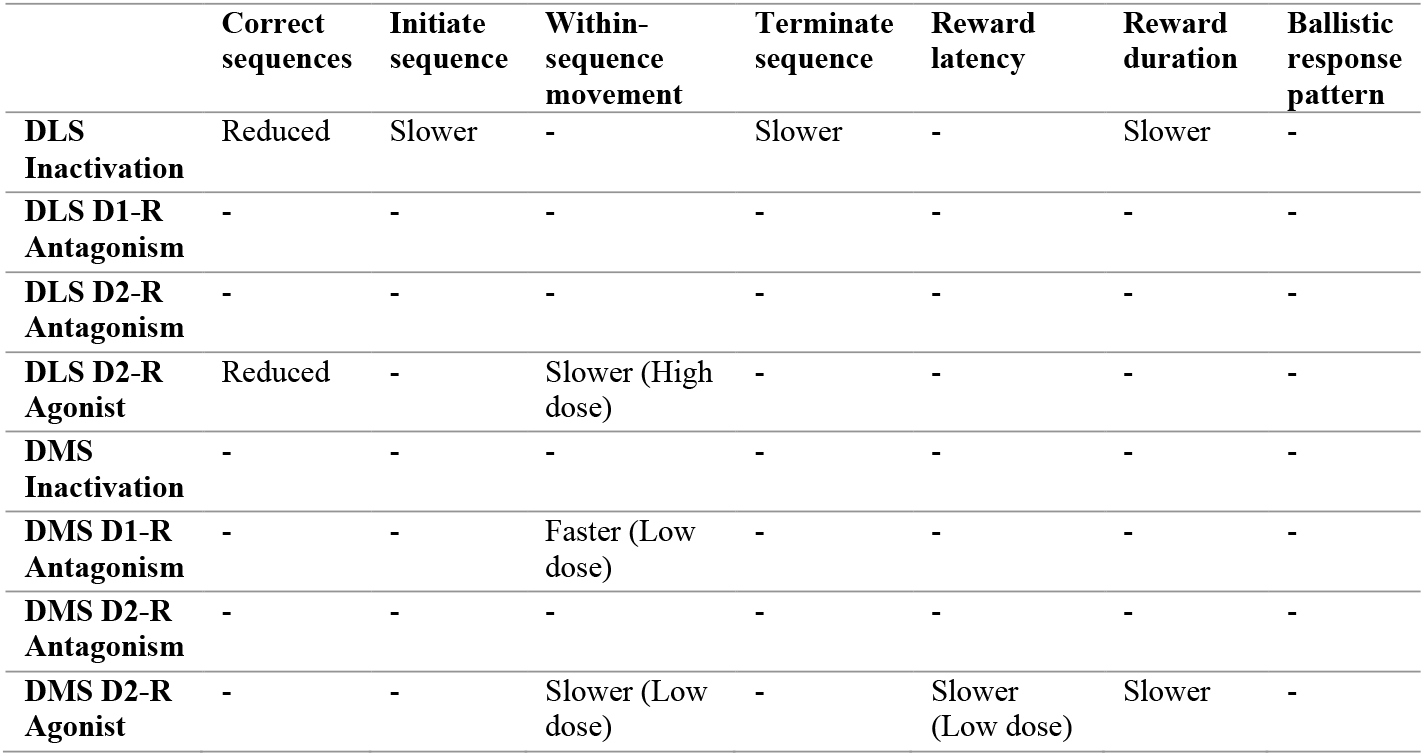
Effect on action sequencing following inactivation and D1-R and D2-R agonism and antagonism in the dorsal striaum.

### Dorsal Striatum Inactivation

DMS inactivation, like previous excitotoxic lesions of the DMS (Turner et al 2022), did not impact action sequencing in expertly performing rats. These results are consistent with a role for the DMS in learning rather than trained performance following extensive training (Dhawale et al. 2021; Kupferschmidt et al. 2017; Thorn et al. 2010; Yin et al. 2005). However, DLS inactivation resulted in rats completing fewer trials and correct sequences with an extended delay to initiate sequences without a significant change in the movement time between nose poke holes or to the magazine, reminiscent of the effects observed following pre-training DLS excitotoxic lesions (Turner et al. 2022). In addition, inactivation resulted in elongation of the terminal sequence response and time spent in the magazine following reward delivery. Collectively these results demonstrate that the DLS plays a specific role in transitioning between action chunks, with delayed execution of initiation and termination elements. These events mark the transitions between the nose poking phase and reward collection phase which appeared to be executed as separate action chunks demarcated by pauses.

Evidence supporting models of sequential or parallel function has shown that DMS activity peaks early in learning then dissipates as DLS activity increases with experience (Bergstrom et al. 2018; Kupferschmidt et al. 2017). Kupferschmidt et al. (2017) observed a reduction in DMS and DLS activity with training, however with different dynamics such that DMS disengagement occurs once the task is learned whereas DLS activity decays more gradually. Based on neural recordings, Thorn et al. (2010) suggested that the DLS is capable of control early in acquisition but does not fully attain control until stable performance is achieved, when DMS activity subsides, thus removing competition imposed via downstream targets. In support of a competitive relationship between the DMS and DLS, it has been shown on a spatial alternation maze task (where flexible responding is optimal) that DMS lesions impaired, while DLS lesions actually improved, task acquisition (Moussa et al. 2011) - one of the first studies to demonstrate that DLS ablation results in faster acquisition of goal-directed learning. Further, examination of a simple conditional stimulus-response rule in marmosets found inactivation of the putamen (homologous to the DLS) impaired performance, while low doses in the caudate (homologous to the DMS) improved performance (Jackson et al. 2019). Similarly, Turner et al. (2022) found acquisition of action sequencing was more rapid following DMS lesions, consistent with the argument above for the DMS constraining DLS functions and further supporting the suggestion that there is some level of a competitive or permissive relationship between the DMS and DLS.

Our results are consistent with this known role of DLS in the performance of skills and habits (Rueda-Orozco and Robbe 2015; Smith and Graybiel 2013; Yin et al. 2004; 2006), and with task bracketing patterns identified from DLS recordings (Barnes et al. 2005; Jin and Costa 2010; Jin et al. 2014; Jog et al. 1999; Kubota et al. 2009; Smith and Graybiel 2013; Thorn et al. 2010). Formal accounts of action chunking suggest chains of actions can become “chunked” such that they are selected and executed as a group (Dezfouli and Balleine 2013; Dezfouli et al. 2014). There are two parallel processes thought to be involved in chunk formation, segmentation that creates break points within long strings of actions, and concatenation that binds elements within these segments together (Fonollosa et al. 2015; Wymbs et al. 2012). Neurons in the dorsal striatum have been shown to produce a start/stop or bracketing pattern around the execution of chunked task elements (Geddes et al. 2018; Jin and Costa 2010; Jin et al. 2014; Jog et al. 1999; Smith and Graybiel 2013; Vandaele et al. 2019). Based on this pattern, the DLS has been suggested to be more important for starting and ending or selecting chunks, rather than representing all the individual sequence components (Markowitz et al. 2018; Rueda-Orozco and Robbe 2015). Jin and Costa (2010) showed that the sequence start/stop pattern emerges with learning and that deletion of striatal NMDA receptors prevented mice from refining action sequence performance, without changes to within-sequence press rate. Further, Cunningham et al. (2021) provided evidence that the DLS task-bracketing pattern reflects what occurred on the previous trial more consistently than the current trial. This raises the possibility that DLS task bracketing patterns reflect the selection of a prior sequence for repetition. Our results provide further evidence that DLS activity is important for action selection and initiation, rather than the individual motor actions *within* the chunk, given the effects on initiation and termination without changes to nose poke behaviour within the sequence.

In a previous study exploring freely moving self-initiated movement sequences, Markowitz et al. (2018) showed that DLS lesions caused transitions between movements to became more random – essentially altered ‘grammar’ - without changes to the underlying individual behaviours. They suggest the DLS is required to assemble sequences of actions and/or the transitions between actions, but not for executing the individual actions themselves (Markowitz et al. 2018). However, this may depend on whether the behaviour is learned or innate. Dhawale et al. (2021) address predictions from the action selection and vigour modulation models yet propose an additional control function. They show likelihood of learned actions is dependent on the DLS, however DLS manipulations lead to the same but slower actions, indicating an additional role in response vigour. Movement trajectories also reflected early learning patterns suggesting a role in fine-grained kinematics of learned behaviour. A key difference they point out is that innate or naturally expressed behaviours may be more DLS-independent (e.g. grooming), but that learned, skilled or dynamically corrected actions may require DLS for both learning and expression. However, this was based on investigation of a singular action, so it is difficult to determine if this response slowing is related to initiation, execution or termination processes.

While Markowitz et al. (2018) developed this idea from innate exploratory movements, such as turn left or run, and Dhawale et al. (2021) used a timed lever press. Here we expand this to a more complex sequence of five uncued instrumental responses that must be executed in the correct order to receive a reward. Longer sequences such as this are required to allow separation of initiation, execution and termination elements. However, longer sequences are at risk of not becoming automatic or breaking into smaller chunks. The ballistic response pattern, as well as the pauses before and after each sequence observed in this study, is consistent with the formation of chunked action sequences. Further, the pause patterns observed here suggest the nose poke sequence chunk is separable from the chunk of actions required to collect the reward. This contrasts with the suggestion that the striatum is responsible for the concatenation of chunk elements, and instead suggests a role for the DLS in transitioning between chunks. Perhaps this role is not to control individual actions, but to control the selection of the next sequence from a repository of previously trained actions that have been elicited within the current environment (e.g., context dependent). However, this process may lack adaptability when the current environment is not represented within the repository or if there are subtle changes that have not been detected by the DLS (e.g. change in outcome value).

### Dopamine D1/D2 receptors

The action of dopamine at D1 receptors is restricted to post-synaptic sites, however the diverse expression and actions of D2 receptors is more complex. D2 receptors are primarily located postsynaptically where dopamine and D2-R agonists *reduce* excitability (Hernandez-Lopez et al. 2000). Striatal D2 receptors are also found on cholinergic interneurons (CINs) where activation reduces firing, leading to disinhibition of connected SPNs and local modulation of dopamine release (Gallo 2019). D2 receptors are also expressed presynatically as autoreceptors on the terminals of incoming dopaminergic neurons with a role in regulating the synthesis, release and re-uptake of dopamine (Ford 2014). Further, D2 autoreceptors are expressed on cortical inputs where activation inhibits corticostriatal activity (Bamford et al. 2004). This configurational complexity is reflected in the often biphasic or opposing response to pharmacological manipulations of D2 receptors (Eilam and Szechtman 1989; Van Hartesveldt et al. 1992). Early studies showed extracellular DA levels drop following striatal infusion of the D2-R agonist quinpirole, likely due to autoreceptor regulation of release and re-uptake (Kurata and Shibata 1990). While subsequent studies using specific D2 knock-out models and local manipulations suggest that D2 receptors on SPNs play a more significant role than those on CINs, at least for motor control (Gallo 2019). Whether these non-linear findings from locomotion studies can be extrapolated to other behaviours, such as motivation, learning and skilled actions, is yet to be determined however our results suggest biphasic dose-dependent effects on action sequences.

In this study, the D2-R *agonist* quinpirole resulted in several changes across both regions. Firstly, in the DMS, the low dose led to significantly longer intervals between nose pokes, longer reward collection latency, and longer time spent in the magazine. The cumulative effect being longer sequence duration. In contrast, the high dose of quinpirole in the DMS only increased the time spent in the magazine. This is likely due to the opposing dose-dependent effects of pre- and post-synaptic D2-R activation, whereby low doses act presynatically on high affinity autoreceptors reducing DA release, however at higher doses quinpirole acts on low affinity postsynaptic D2 receptors mimicking DA release (Horst et al. 2019). Importantly, these doses have been shown to not induce locomotor changes beyond the first 10 minutes following infusion in the dorsal striatum (Van Hartesveldt et al. 1992). In this context, our results show that *decreasing* DMS indirect pathway activity specifically delays within-chunk sequence movements and reward related actions. However, *increasing* DMS indirect pathway activity only increased time to consume the reward, again providing evidence that this circuit is not critical to sequence selection, but DMS perturbations can impact reward-related behaviours, consistent with a role in incorporating outcome value and contingency in decision-making processes.

In the DLS, low dose quinpirole specifically increased the number of incorrect sequences, supporting the proposal that DLS D2-R activation is important for inhibiting alternative actions (Vicente et al. 2016). By activating D2 autoreceptors and reducing DA release, alternative actions may become less constrained resulting in the observed increase in incorrect responses. In contrast, high dose quinpirole, which would act postsynaptically, reduced the number of correct sequences and increased the latency between nose pokes with a trend towards increased sequence duration. The inhibitory D2-indirect pathway is suggested to be important for supressing alternative responses from being selected. Given high doses increase circuit inhibition, excessive inhibition may lead to excessive inhibition of actions and a subsequent reduction in total correct sequences, without an increase in incorrect sequences and longer intervals between actions. There was however no effect of D2 receptor antagonism, suggesting blocking postsynaptic D2 receptors is not sufficient to alter behaviour. Collectively, these results suggest greater sensitivity to artificially increasing D2 auto- and postsynaptic-receptor activity as opposed to decreasing D2-R activation, consistent with the proposed role of iSPNs in selective prevention of alternative actions.

Infusion of the D1-R antagonist SCH23390 or the D2-R antagonist raclopride in the DMS and DLS left performance intact, except for faster movement between nose poke holes following DMS infusion of the low dose SCH23390. Interestingly, this acceleration was only detected within the five-step sequence and not for other movements, suggesting that although DMS inactivation was non-consequential, altering D1-R activation impacts within-sequence execution. This is consistent with our prior findings that sequencing is more efficient following DMS loss of function and suggests rapid sequence performance may be constrained by activity of the DMS D1 direct pathway.

The lack of effect follow D1-R antagonist infusion is consistent with a role in learning rather that habits with others finding a similar lack of effect in the DLS alongside effects in the nucleus accumbens (Verharen et al. 2019). There was also no effect of the D2-R antagonist, indicating activation of the indirect pathway from either the DMS or DLS does not alter sequence performance. This result was unexpected given DLS infusions of raclopride within the dose range used in the current study have been shown to restore habitual responding back to goal-directed control (Corbit et al. 2014) and impair reversal learning (Sala-Bayo et al. 2020). Raclopride has also been shown to delay lever press reaction times (Keeler et al. 2014), albeit at slightly higher doses. It has been suggested the inhibitory influence of D2-R activation may help to shape or select actions (Keeler et al. 2014). Therefore, blocking the indirect pathway may be sufficient to unlock adaptive responding following devaluation, but not alter overtrained action sequences where alternative responses would not be adaptive.

The use of local pharmacological infusions has some limitations. Drug infusions do not provide the temporal precision required to decipher how specific activity patterns within and between systems may contribute to sequence performance as has been examined using optogenetic tools (Tecuapetla et al. 2016). It is important to note that while GABA-A/B agonists muscimol and baclofen have been commonly used to inactivate the dorsal striatum (Corbit and Janak 2007; Corbit et al. 2012; Jackson et al. 2019; Vandaele et al. 2019; Yin et al. 2006), the mechanism of action is complicated as both interneurons and SPNs are GABAergic. Further, the mechanism of action of many agents is not completely isolated to the target receptor and is not restricted to a specific cell type. For example, the actions of quinpirole are likely to impact D2 presynaptic autoreceptors, postsynaptic receptors as well as D2 receptors on cholinergic interneurons in the vicinity. However, this would also be the case with native dopamine transients, hence these results provide insight into the heterogenous role of the D2 system in the dorsal striatum more broadly. The use of different tasks is likely to be critical to our understanding of action control and skilled performance. Each has limitations in terms of steps, transitions, spatial and temporal dynamics but collectively should help to build a more robust model of striatal function. In everyday routines, we frequently move between different acquired chunked sequences and these transitions from one sequence to the next may be disrupted in compulsive disorders. Therefore, we need experimental approaches that allow us to investigate movement between sequences to better understand human behaviour. The results of this study highlight the importance of incorporating findings from more complex designs in our understanding of neural control over behaviour, however we still have much work to do in order to understand complex behavioural repertoires.

## Conclusions

Overall, these results demonstrate that performance of learned action sequences requires DLS, but not DMS, and involve optimal states of D2, but not D1 receptors, highlighting a role for the indirect pathway in sequence selection and execution. These results provide new insights into the how neural circuits support the execution of skilled motor actions. They also highlight the importance of using tasks where it is possible to isolate initiation, execution and termination components of action sequences to determine if regions are required to sequence individual elements or are required for the initiation and execution of such sequences.

## Acknowledgements

We would like to thank Prof. Amy Milton for supporting this research under her PPL Home Office Licence.

